# The genome of *Auriculariopsis ampla* sheds light on fruiting body development and wood-decay of bark-inhabiting fungi

**DOI:** 10.1101/550103

**Authors:** Éva Almási, Neha Sahu, Krisztina Krizsán, Balázs Bálint, Gábor M. Kovács, Brigitta Kiss, Judit Cseklye, Elodie Drula, Bernard Henrissat, István Nagy, Mansi Chovatia, Catherine Adam, Kurt LaButti, Anna Lipzen, Robert Riley, Igor V. Grigoriev, László G. Nagy

## Abstract

The Agaricomycetes are fruiting body forming fungi that produce some of the most efficient enzyme systems to degrade woody plant materials. Despite decades-long interest in the ecological and functional diversity of wood-decay types and in fruiting body development, the evolution of the genetic repertoires of both traits are incompletely known. Here, we sequenced and analyzed the genome of *Auriculariopsis ampla*, a close relative of the model species *Schizophyllum commune*. Comparative analyses of wood-decay genes in these and other 29 Agaricomycetes species revealed that the gene family composition of *A. ampla* and *S. commune* are transitional between that of white rot species and less efficient wood-degraders (brown rot, ectomycorrhizal). Rich repertoires of suberinase and tannase genes were found in both species, with tannases generally restricted to species that preferentially colonize bark-covered wood. Analyses of fruiting body transcriptomes in both *A. ampla* and *S. commune* highlighted a high rate of divergence of developmental gene expression. Several genes with conserved developmental expression were found, nevertheless, including 9 new transcription factors as well as small secreted proteins, some of which may serve as fruiting body-specific effector molecules. Taken together, the genome sequence and developmental transcriptome of *Auriculariopsis ampla* has highlighted novel aspects of wood-decay diversity and of fruiting body development in mushroom-forming fungi.

## Introduction

Mushroom-forming fungi (Agaricomycetes) are of great interest for comparative genomics due to their importance as wood-degraders in global carbon cycling and as complex multicellular organisms that produce agriculturally or medicinally relevant fruiting bodies. Recent advances in genome sequencing has brought new light on several aspects of lignocellulose decomposition and the genetic repertoire of fruiting body development in mushroom-forming fungi ^1–5^.

The Agaricomycetes display diverse strategies to utilize lignocellulosic substrates. While genomic analyses have helped to uncover the main patterns of duplication and loss of plant cell wall degrading enzyme (PCWDE) families, our understanding of the enzymatic repertoires of Agaricomycetes and how they use it to degrade various lignocellulosic components of plants is far from complete. Fungi have traditionally been classified either as white rot, in which all components of the plant cell wall are being degraded ^6^, or brown rot, in which mostly cellulosic components are degraded, but lignin is left unmodified or only slightly modified ^7^. Several species have been recalcitrant to such classification, which prompted a reconsideration of the boundaries of the classic WR and BR dichotomy ^8,9^. Such species are found across the fungal phylogeny, but seem to be particularly common among early-diverging Agaricales, including the Schizophyllaceae, Fistulinaceae and Physalacriaceae ^2,8,9^. *Schizophyllum commune*, the only hitherto genome-sequenced species in the Schizophyllaceae, for example, produces white rot like symptoms, but lacks hallmark gene families (e.g. lignin-degrading peroxidases) of WR fungi ^2,8–10^. Accordingly, it lacks the ability to degrade lignin and achieves weak degradation of wood ^9,11,12^, although this might be complemented by pathogenic potentials on living plants or the activity of other, more efficient degraders that co-inhabit the same substrate^13^. Analyses of the secretome and wood-decay progression of *S. commune* revealed both WR and BR-like behaviors ^10,14^, although several questions on the biology of this species remain open.

Fruiting body production, is a highly integrated developmental process triggered by a changing environment, such as a drop in temperature, nutrient depletion or shifts in light conditions ^15–17^. It results from the concerted expression of structural and regulatory ^1,18–22^ genes as well as other processes, such as alternative splicing ^3,23^, allele-specific gene expression ^3^ and probably selective protein modification ^3,24^. Known structural genes include hydrophobins ^25–27^, lectins ^28–30^, several cell wall chitin and glucan-active CAZymes ^5,31–33^, and probably cerato-platanins, expansin-like ^3,4^ and an array of other genes ^34^. Regulators of fruiting body development have been characterized in several species, in particular in *Coprinopsis cinerea* ^18,19,35–37^ and *S. commune* ^1,2,24^. Despite much advance in this field, several aspects of fruiting body development are quite poorly known, including, for example what genes have conserved developmental roles across fruiting body forming fungi or how cell-cell communication is orchestrated in developing fruiting bodies. *S. commune* has served as a model organism for fruiting body development for a long time ^2,15,16^. This species, like other Schizophyllaceae (e.g. the genus *Auriculariopsis*) produce cyphelloid fruiting bodies, which are reduced morphologies derived from more complex ancestors. Cyphelloid fruiting bodies are inverted cup-like forms with unstructured (e.g. *A. ampla*) or slightly structured (e.g. *S. commune*) spore-bearing surfaces (hymenophore). Albeit the hymenophore structure of *S. commune* resembles gills (hence the common name ‘split gill’), it is not homologous to real gills of mushrooms, rather, it results from the congregation of several individual cup-like fruiting bodies.

We sequenced the genome of *Auriculariopsis ampla*, a close relative of *S. commune* that primarily inhabits the bark of dead logs and produces simple, cup-shaped fruiting bodies. Through analyses of gene repertoires for plant cell wall degradation in *A. ampla, S. commune* and 29 other Agaricomycetes, we detect signatures of adaptation to wood colonization through the bark and suggest that these two species have unusual plant cell wall degrading enzyme repertoires. By sequencing developmental transcriptomes of *A. ampla* and comparing it to that of *S. commune*, we identify conserved developmental genes that might be linked to fruiting body development, including small secreted proteins, some of which show extreme expression dynamics in fruiting bodies.

## Results and Discussion

### *Auriculariopsis* has a typical Agaricales genome

The genome of *Auriculariopsis* was sequenced using PacBio, assembled to 49.8 Mb of DNA sequence in 351 scaffolds with a mean coverage of 54.38x (343 scaffolds >2 kpb, N50: 19, L50: 0.53 Mb). We predicted 15,576 protein coding genes, based on which BUSCO analysis showed a 98.6% degree of completeness (273 complete, 28 duplicated, 2 fragmented, 2 missing). We included *A. ampla* and its close relative *Schizophyllum commune* in a comparative analysis with 29 other Agaricomycetes. A species phylogeny was reconstructed from 362 single-copy orthologs (14,2436 amino acid characters) for the 31 taxa; the inferred topology resembles published genome-scale trees of Agaricomycetes very closely and received strong (>85%) bootstrap support for all but two nodes (Figure 1a). *Auriculariopsis ampla* clustered with *S. commune*, together representing the Schizophyllaceae, with their immediate neighbor being *Fistulina hepatica* (Fistulinaceae). The gene repertoire of *A. ampla* (15,576 genes) is very similar to that of *S. commune* ^2^(16,319 genes) and the average gene count in the analyzed Agaricales species (17,655), but more than that of *F. hepatica* ^9^ (11,244 genes). We found 8 significantly overrepresented (p-value<=0.05) and 16 underrepresented (p-value<=0.05) InterPro domains in both species, relative to the other 29 species (Supplementary Table 1).

**Figure 1.**
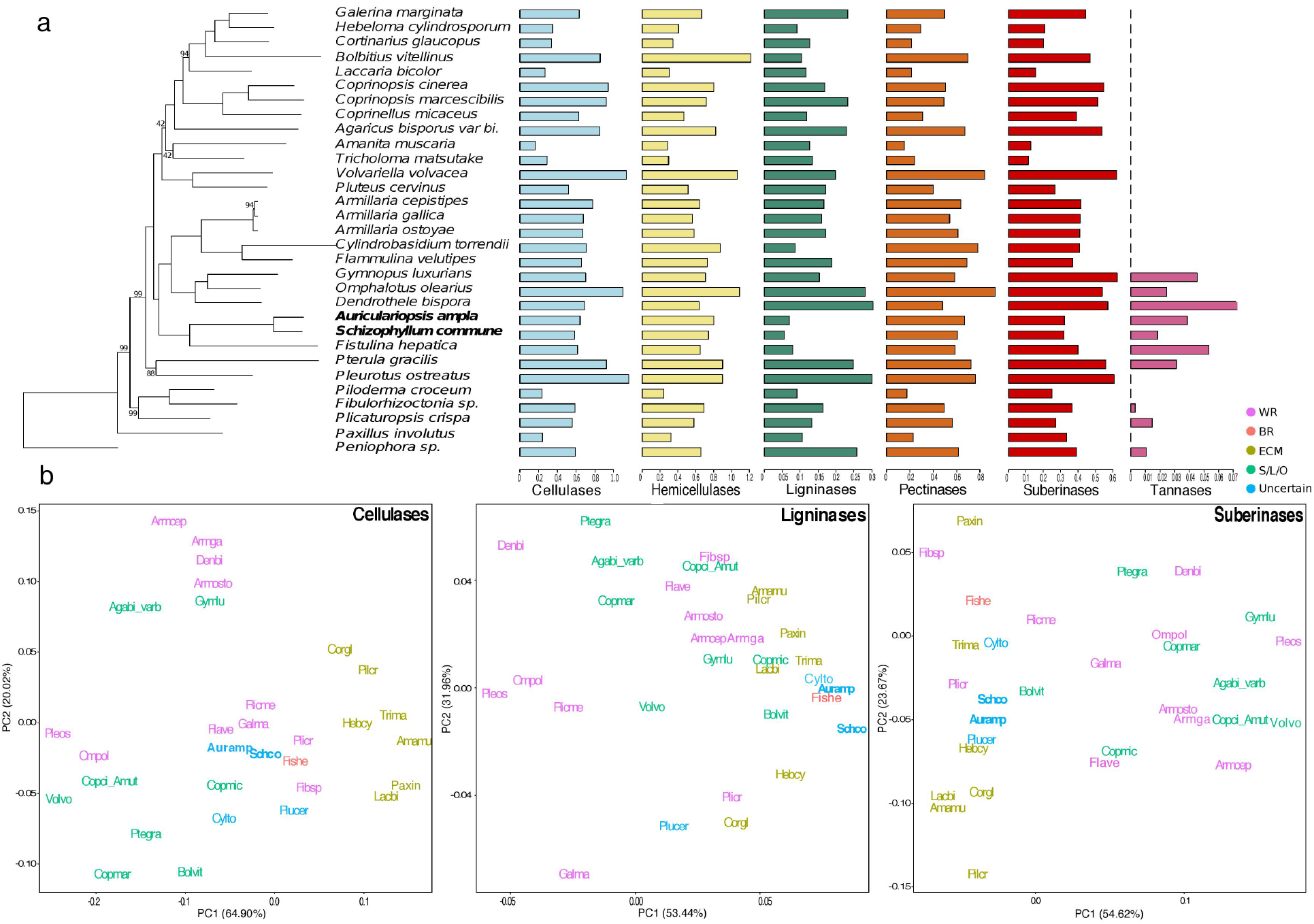
Phylogenetic relationships and lignocellulose degrading gene repertoire of *A. ampla* compared to *S. commune* and 29 other Agaricomycetes. A, species tree showing the phylogenetic affinities of the Schizophyllaceae (bold, left panel) and copy number distribution of cellulose, hemicellulose, pectin, lignin degrading gene families as well as those of putative suberinases and tannases. B, phylogenetic principal component analyses of cellulose, lignin and suberin degrading enzymes. Species names colored based on nutritional mode (WR - white rot, BR - brown rot, ECM - ectomycorrhizal, S/L/O - soil and litter decomposer, Uncertain - nutritional mode not known with certainty). For better visibility, a few species have been moved slightly on the plots (information in Supplementary Table 5) See also Supplementary Figure 1 for original plots.

### *Auriculariopsis* and *Schizophyllum* have unusual wood-decay strategies

The substrate-wise phylogenetic PCA portrays a separation of these two species from the other 29 species used in this dataset. In the case of cellulases, these two species cluster together with WRs and S/L/O, suggesting a similar arsenal of CAZymes acting on the cellulose degradation (Figure 1b). Enzyme families acting on crystalline cellulose (cellobiohydrolases - GH6, GH7) were present in lower numbers than in WRs and S/L/O species, but similar to ectomycorrhizal ones. The pattern was mostly identical for hemicellulases and pectinases (Figure 1b) where CAZyme copy numbers were similar to that of white rotters and litter decomposers. However, some CAZymes related to xylanase and pectinase activities including xylosidases, pectate lyases, pectin acetylesterases, and acetyl xylan esterases (AA8, GH30, GH43, GH95, CE12, PL1, PL3, PL4) have higher copy numbers in the two species than in most WRs. This could imply their role in hemicellulose and pectin degradation, as reported previously ^10^. However, ligninolytic CAZymes revealed a clear difference from WR species. The absence of class II peroxidases in both species ^6–8^, made them cluster towards ectomycorrhizal and BR species. Copper radical oxidases (CROs) are known to be responsible for the production hydrogen peroxide and were also reported to have a significantly different repertoires in BRs and WRs ^9^. In our analysis we found very low numbers of CROs (AA5) in *Auriculariopsis* and *Schizophyllum* as compared to ECM, S/L/O and WR species.

Because *A. ampla* and *S. commune* often occur on bark as first colonizers, we also examined protein families that putatively degrade important bark compounds. Suberin, lignin and tannins represent the major components of bark ^38–40^. We built on previous datasets to obtain putative suberinase ^38,39^ and tannase ^40^ copy numbers for 31 species in our dataset. In general, suberin comprises aromatic compounds cross linked by poly-aliphatic and fatty-acid like components which requires extracellular esterases and lipases for their breakdown ^38^. Based on phylogenetic PCA of putative suberinases *A. ampla* and *S. commune* were transitional between typical WR and ECM, BR (*Fistulina*), uncertain (*Cylindrobasidium, Pluteus*) or tentative WR (*Fibulorhizoctonia*) species. This separation is most pronounced along the first axis (PC1), the main contributor of which is the AA3 family. *A. ampla* and *S. commune* had few genes in this family, similar to most ECM species. In terms of most other families, *A. ampla* and *S. commune* resembled WR species. The cutinase (CE5) repertoires of the two species are similar to those of litter decomposers and certain WR taxa (e.g. *Galerina*, *Dendrothele*, *Fibulorhizoctonia*, and *Peniophora*), although this family was missing from several WR species. Tannin acyl hydrolases (tannase, EC 3.1.1.20) are responsible for the degradation of tannins, polyphenolic plant secondary metabolites characteristic to the bark and wood tissues. Tannases were found in 10 out of 31 species, mostly in those that occur preferentially on bark, such as *Auriculariopsis*, *Schizophyllum*, *Peniophora*, *Dendrothele* and *Plicaturopsis*, and a few others (*Gymnopus*, *Pterula*, *Fibulorhizoctonia*, *Omphalotus* and *Fistulina*). This could indicate a specialization of these species to substrates with high tannin content, such as bark, suggesting adaptations to the early colonization of bark-covered wood. Notably, *Pluteus*, a species with an uncertain nutritional mode, groups closely together with *A. ampla* and *S. commune* on the suberinase PCA, although it had low pectinase, hemicellulase and cellulase copy numbers, leading to a position close to ECM species and some litter decomposers in other PCA analyses (Supplementary Figure 1).

### Transcriptomics reveals a high rate of developmental evolution

*Auriculariopsis ampla* and *S. commune* have a similar developmental progression (Figure 2a-2e), permitting a comparison of their transcriptional programs. Fruiting body development starts in both species with the appearance of minute globose primordia (stage 1 primordia), in which a cavity develops (Stage 2 primordia). This cavity further expands in *A. ampla* to produce an open, pendant fruiting body (Young fruiting body and fruiting body stages), whereas in *S. commune* several such units form a multi-lobed assemblage.

**Figure 2.**
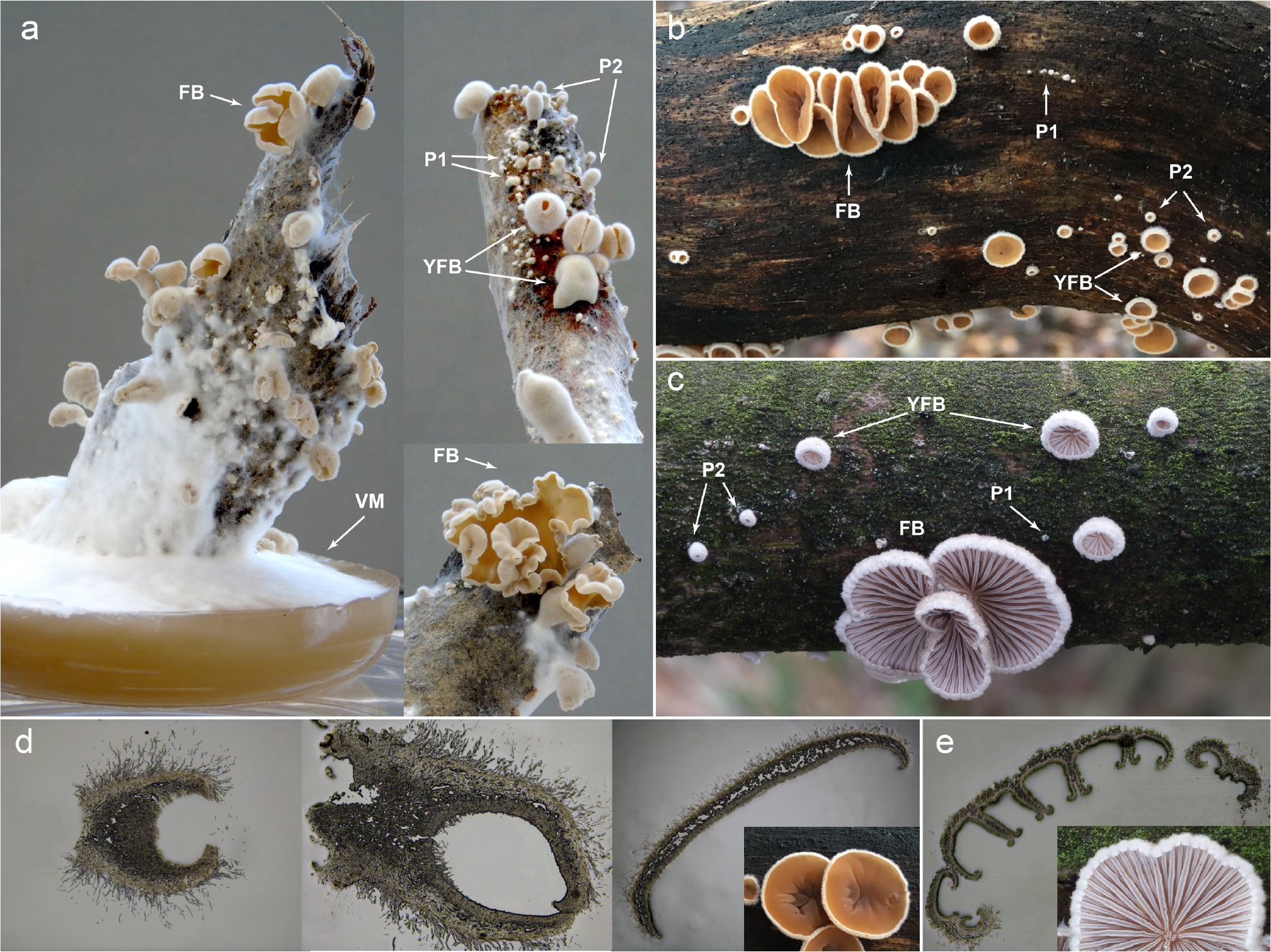
Fruiting bodies and developmental stages of *A. ampla* and *S. commune*. Developmental stages are indicated on each panel. A, fruiting bodies of *A. ampla* produced in vitro, on sections of barked poplar logs plugged into malt-extract agar. B and C, fruiting bodies of *A. ampla* and *S. commune* in their natural habitat. D, cross sections of developmental stages of *A. ampla*: left - stage 1 primordium (left), stage 2 primordium (middle) and mature fruiting body (right). E, Cross section of a mature fruiting body of *S. commune*, showing congregated single fruiting bodies.

For comparison with *S. commune*, we generated RNA-Seq data from 5 developmental stages of *A. ampla* (vegetative mycelium, stage 1 and stage 2 primordia, young and mature fruiting bodies, see Figure 2a-b,d) in biological triplicates, >30 million (30-78M, mean: 46M) paired-end 150 base reads for each sample on Illumina platform (mean read mapping: 83%, Supplementary Table 2). Corresponding data for the same developmental stages (Figure 2c, e) for *S. commune* were taken from (ref. 3). Based on global transcriptome similarity, fruiting body samples grouped together away from vegetative mycelium ones (Figure 3a) consistent with the complex multicellular nature of fruiting bodies as opposed to a simpler cellularity level of vegetative mycelia. Among the fruiting body samples, stage 1 and stage 2 primordia were similar to each other, whereas young fruiting bodies and mature fruiting bodies formed distinct groups. We identified 1466 developmentally regulated genes in *A. ampla*, which is similar in magnitude to that reported for *S. commune*, but less than that for more complex species (e.g. *Coprinopsis*, *Armillaria*; taken from (ref. 3). Of the developmentally regulated genes, 967 showed a significant (≥4) fold change in the transition from vegetative mycelium to stage 1 primordia. In terms of significantly differentially expressed genes (DEGs), the highest numbers of up and downregulated were also found between vegetative mycelium and stage 1 primordia (1166 and 842 genes, Figure 3b), which is consistent with the position of samples on the MDS plot (Figure 3a). Much fewer genes were differentially expressed between stage 1 and 2 primordia and between stage 2 primordia and young fruiting bodies. In fruiting bodies, we found 110 and 37 significantly up- and downregulated genes, respectively, a comparatively higher number that is potentially related to sporulation.

**Figure 3.**
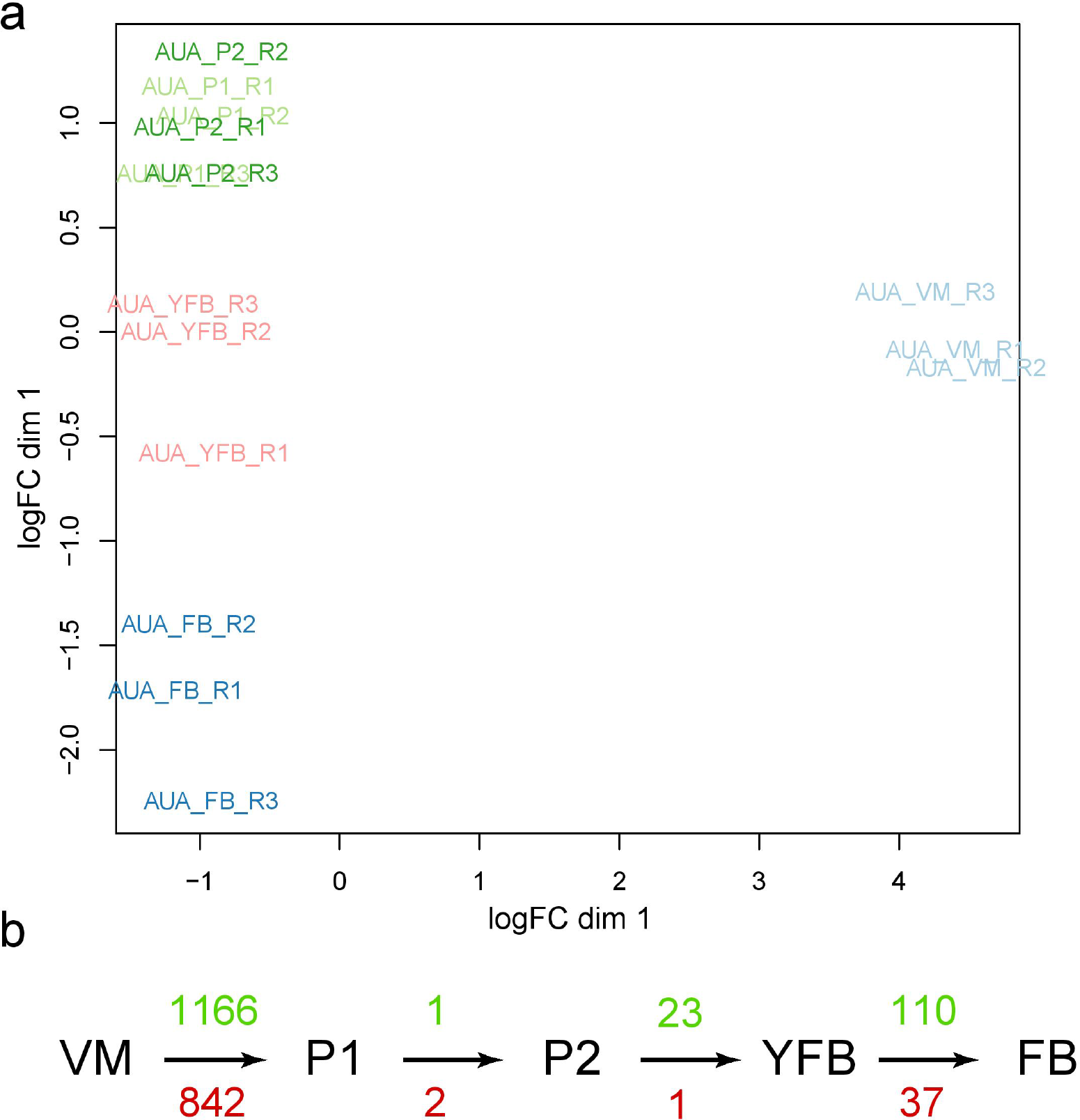
Overview of the developmental transcriptome of *A. ampla*. A, Multi-dimensional scaling for RNA-Seq replicates from 5 developmental stages of *Auriculariopsis ampla*. Biological replicates belonging to similar tissue type group together. The replicates for P1 and P2 cluster together and remaining developmental stages keep apart. B, Graphical representation of number of significantly upregulated (green) and downregulated (red) genes among developmental stages and tissue types in *A. ampla*. Abbreviations: VM - vegetative mycelium, P1 - stage 1 primordium, P2 - stage 2 primordium, YFB - young fruiting body, FB - mature fruiting body.

We assessed the similarity between the 2 species’ developmental transcriptomes by analyzing the expression of one-to-one orthologous gene pairs, hereafter referred to as co-orthologs. To identify co-orthologs, proteomes of *A. ampla* and *S. commune* were clustered into 18,804 orthogroups using MCL, of which 7463 represented co-orthologs. Of these, 7369 co-orthologs were expressed under our experimental conditions in both species (Supplementary Table 3). Pairwise comparison between developmental stages showed highest similarity within species across all 7369 co-orthologs (Figure 4a). This pattern was more pronounced in an analysis of developmentally regulated co-orthologs (Figure 4b, see Methods), indicating that developmental gene expression in *A. ampla* and *S. commune* has diverged since their common ancestor so that similarity between homologous fruiting body stages of the two species is lower than that between different stages of the same species. Vegetative mycelia of both species differed most from all other stages of the same species but showed some similarity across species. Similarly, we observed a strong similarity between young fruiting bodies and fruiting bodies of *A. ampla* and *S. commune*, indicating that late stages of fruiting body development share more similarity across species than do early stages. Similar patterns were observed when the analyses were restricted to co-orthologous transcription factors (Figure 4c) and its developmentally regulated subset (Figure 4d). Similarity between late developmental stages of the two species was more pronounced in the analysis of developmentally regulated genes. Given that *A. ampla* and *S. commune* are each other’s closest relatives, the low similarity of gene expression among their fruiting bodies (Figure 4) indicates that developmental gene expression has diverged at a high speed since their divergence. This is surprising in comparison to similar analyses of animals, where gene expression patterns could be predicted from tissue identities across the entire mammalian clade ^41^. This suggests that fruiting body development evolved at a high rate, erasing identities of similar developmental stages across species. Nevertheless, these data show that during fruiting body maturation gene expression dynamics shows conserved patterns among phylogenetically closely related species. These data further indicate that there should be genes with similar expression profiles in *A. ampla* and *S. commune*.

**Figure 4.**
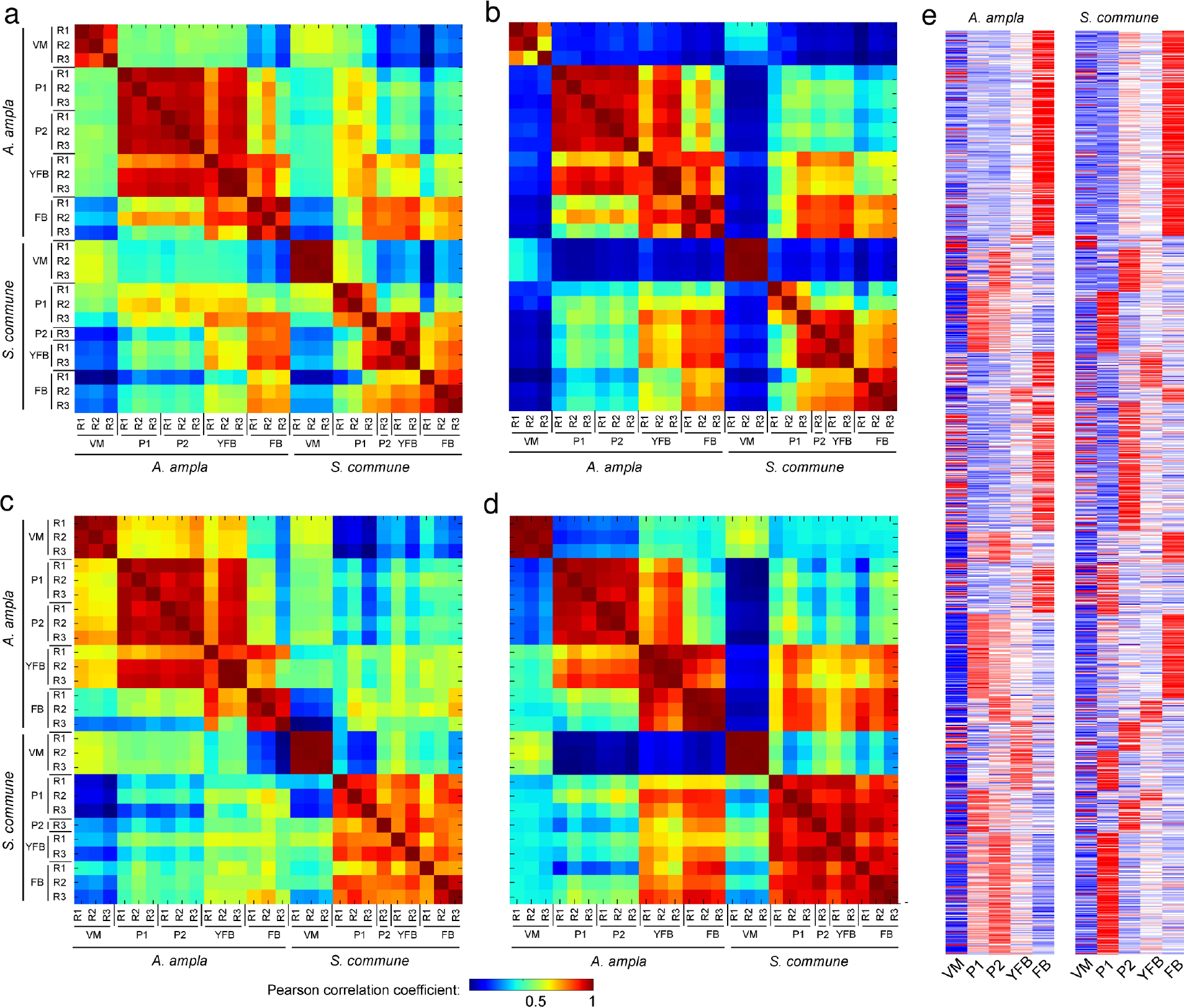
Global transcriptome similarity between developmental transcriptomes of *A. ampla* and *S. commune*. Pearson correlation coefficient-based heatmaps show similarity among developmental stages of the two species for all 7369 co-orthologs (A), for 1182 developmentally regulated co-orthologs (B), for 252 co-orthologous transcription factor pairs (C) and 42 developmentally regulated co-orthologous TF pairs (D). Warmer color indicates higher similarity. Biological replicates are indicated next to the heatmap (R1-R3). E, paired heatmap of gene expression (FPKM) for 7369 co-orthologous gene pairs between *A. ampla* and *S. commune*. Developmental stages for both species are as follows: VM - vegetative mycelium, P1 - stage 1 primordium, P2 - stage 2 primordium, YFB - young fruiting body, FB - mature fruiting body.

Despite the low global similarity, several genes with conserved expression patterns could be identified. The most highly upregulated co-ortholog in *A. ampla* and *S. commune* was a heat shock protein 9/12 family member, that is homologous to *Aspergillus nidulans* awh11 and *S. cerevisiae* hsp12, farnesol-responsive heat shock proteins. These genes had a significant upregulation in stage 1 primordia of both *A. ampla* and *S. commune* (fold change 254x and 855x, respectively) and had high expression values in all fruiting body tissues (>5,000 FPKM, maximum fold change within fruiting bodies 3.4-3.7), suggesting an important role of heat shock proteins during fruiting body development. In further support of this hypothesis, homologs of these genes were found developmentally regulated or differentially expressed also in *Laccaria bicolor* ^21^, *Lentinula edodes* ^42^, *Armillaria ostoyae*, *Coprinopsis cinerea*, *Lentinus tigrinus*, and *Rickenella mellea* ^3^. Another co-orthogroup with large upregulation in stage 1 primordia included A1 aspartic proteases, although the expression dynamics were somewhat different in the two species. We observed an induction in stage 1 primordia in both, but, while upregulation in *A. ampla* was >200x compared to VM, it was only 14x in *S. commune*. Aspartic proteases of the diverse A1 family have been reported as highly induced in fruiting bodies in several previous studies ^3,21,42–44^, although no mechanistic hypothesis for their role in fruiting body development has been proposed yet.

### Putative fruiting body genes are developmentally expressed

We further examined the expression patterns of previously reported fruiting body genes in *A. ampla*. Of the fungal cell wall (FCW) associated genes, hydrophobins were mostly developmentally regulated (8 out of 11 genes) in *A. ampla* (Supplementary Figure 2), often with significantly increased expression coincident with the transition from vegetative mycelium to stage 1 primordia (in six genes), as observed previously ^1,42,45–47^. Several members of two functionally similar families, cerato-platanins (4 of 5 genes dev. reg.) and expansin-like genes (10 of 21 genes) were likewise developmentally regulated. Although cerato-platanins and expansins were often associated with the plant cell wall ^48,49^, their dynamic expression in fruiting bodies suggest potential FCW-related roles. Functional annotations uncovered several putatively FCW-active CAZymes (Supplementary Figure 3), including chitin- and glucan-active GH and GT families, carbohydrate-binding modules, carbohydrate esterases, AA1 multicopper oxidases, AA9 lytic polysaccharide monooxygenases, but also starch synthesizing glycosyl transferases (e.g. GH15, CBM20), which might be related to the mobilization of glycogen reserves during development. Two out of 10 members of the Kre9/Knh1 family were developmentally regulated. This family is involved in *β-*1,6-glucan synthesis and remodeling in *Aspergillus fumigatus* ^50^, *Candida albicans* ^51^, *Saccharomyces cerevisiae* ^52^ and *Ustilago maydis* ^53^ and has been shown to be developmentally expressed in Agaricomycetes fruiting bodies ^3,54^. Its widespread FCW-associated role in both Asco- and Basidiomycota suggests a plesiomorphic role in *β*-glucan assembly in the cell wall and suggests that this family has been co-opted for fruiting body development in Agaricomycetes. Several other previously reported putatively FCW-active CAZyme families ^3,21,42,55–58^ (e.g GH5, GH142 ^5,59–61^, Supplementary Figure 2), also showed developmental expression in *A. ampla*, reinforcing the view that cell wall remodeling is a fundamental mechanism in fruiting body development ^2,3,5,21,31,58,62–64^.

Defense related genes have been in the focus of research on fruiting bodies. We found developmentally regulation of a diverse array of putative defense-related genes by searching for homologs of *Coprinopsis* defense genes ^65^. *A. ampla* and *S. commune* have reduced repertoires of defense-related genes compared to *Coprinopsis cinerea* (Supplementary Figure 2). For example, no homologs of aegerolysins or the ETX/MTX2 pore forming toxin family have been found in their genomes, whereas lectins are only represented by 14 genes as opposed to 39 and 25 in *C. cinerea* and *A. ostoyae*. They have several genes in the thaumatin family, with has been associated with defense ^65^ in both fungi and plants ^66,67^, fungal pathogenicity ^68^ but also with FCW remodeling ^62,69^, depending on the scope of the study. Based on its endo-*β*-1,3-glucanase activity, its efficiency in degrading cell wall components of *Lentinula edodes* ^62^ and *Saccharomyces* ^69^ and developmental expression in axenic fruiting bodies, it appears likely that members of this family are involved in FCW remodeling, although antimicrobial activities have also been predicted for certain members ^3^. Cerato-platanins represent a similar case. This is a family widely expressed in pathogenic fungal-plant interactions ^70,71^, fruiting bodies ^3,4,71^ and defense assays ^65^ and significantly enriched in Agaricomycetes genomes ^3,70^. We detected four developmentally regulated cerato-platanin genes in *A. ampla*, three of which showed an induction at the transition from vegetative mycelium to stage 1 primordia (Supplementary Figure 2). *S. commune* had three developmentally regulated cerato-platanins, with non-matching expression profiles. Further, in *A. ampla* we found 3 developmentally regulated lectin genes (Supplementary Figure 2), as opposed to *S. commune*, which had 8 ^3^. All three genes belong to the ricin-B lectin family and harbor a CBM13 domain, which has demonstrated mannose, N-acetylgalactosamine and xylane binding activities ^72,73^. Ricin-B lectins have been reported as developmentally expressed in fruiting bodies of all Agaricomycetes tested so far ^3,4,28,42,57,65^, although its functions are less clear. It is the largest family of basidiomycete lectins ^3^ and was shown to be toxic to nematodes ^29,74^, although their diverse carbohydrate-binding abilities (mannose and N-acetylgalactosamine) could confer additional or other functions as well.

F-box and BTB/POZ domain containing proteins have recently been reported as an interesting family with probable functions in fruiting body development and a significant expansion in the Agaricomycetes ^3^. *Auriculariopsis* has 246 F-box protein encoding genes, of which 12 were developmentally regulated in our dataset. Of the 96 BTB/POZ domain-containing proteins 26 were developmentally regulated, including some genes with remarkable expression dynamics during development (e.g. fold change 526x, Auramp1_515369). This is similar to figures reported for *S. commune* ^3^. These domains are involved in protein-protein interactions and have been reported to act as transcriptional repressors ^75^, members of selective proteolysis pathways, and include homologs of yeast *Skp1* ^76^ too. Although very little functional information on these families is available in fungi, their expression dynamics in development and previously reported regulatory roles suggest they could be important players in fruiting body development.

### Conserved patterns of transcription factors expression

We examined expression patterns of transcription factors (TFs) and their similarity between the two species. We identified 433 and 437 TFs in the genomes of *A. ampla* and *S. commune* respectively, of which 252 were co-orthologs. These were distributed across 28 TF families, with C2H2-type Zinc finger and Zn (2)-C6 fungal-type TFs being the most dominant (Supplementary Table 4). Individually, 14,5% and 16% of the *Auriculariopsis* and *Schizophyllum* and 17% of the co-orthologous TFs were developmentally regulated, respectively.

These included 5 of the eight previously characterized TF genes of *S. commune* ^1^: *c2h2, gat1, hom1, tea1* and *fst4* showed significant changes in expression, in most cases at the initiation of fruiting body development, whereas *fst3, bri1* and *hom2* showed more or less flat expression profiles (Figure 5). These expression profiles are consistent with previous RNA-Seq based reports ^24^ in *Schizophyllum* and other species ^65,77–79^, except in *hom1* and *gat1*, which, in our data behaved differently, probably due to the different resolution of developmental stage data. The expression profiles of all eight genes were very similar between *A. ampla* and *S. commune*. Homologs of *Lentinula edodes PriB* ^80,81^ (Auramp1_518770, Schco3_2525437) were also developmentally regulated, with an expression peak in stage 1 primordia. Homologues of *Coprinopsis exp1*, which was reported to be involved in cap expansion ^82^, were present in both species and had a matching expression profile, but were not developmentally regulated in our data. In our experiments, *exp1* homologs (Auramp1_481073, Schco3_2623333) showed highest expression in vegetative mycelia and lower expression afterwards, which might be related to the lack of proper caps in *A. ampla* and *S. commune*.

**Figure 5.**
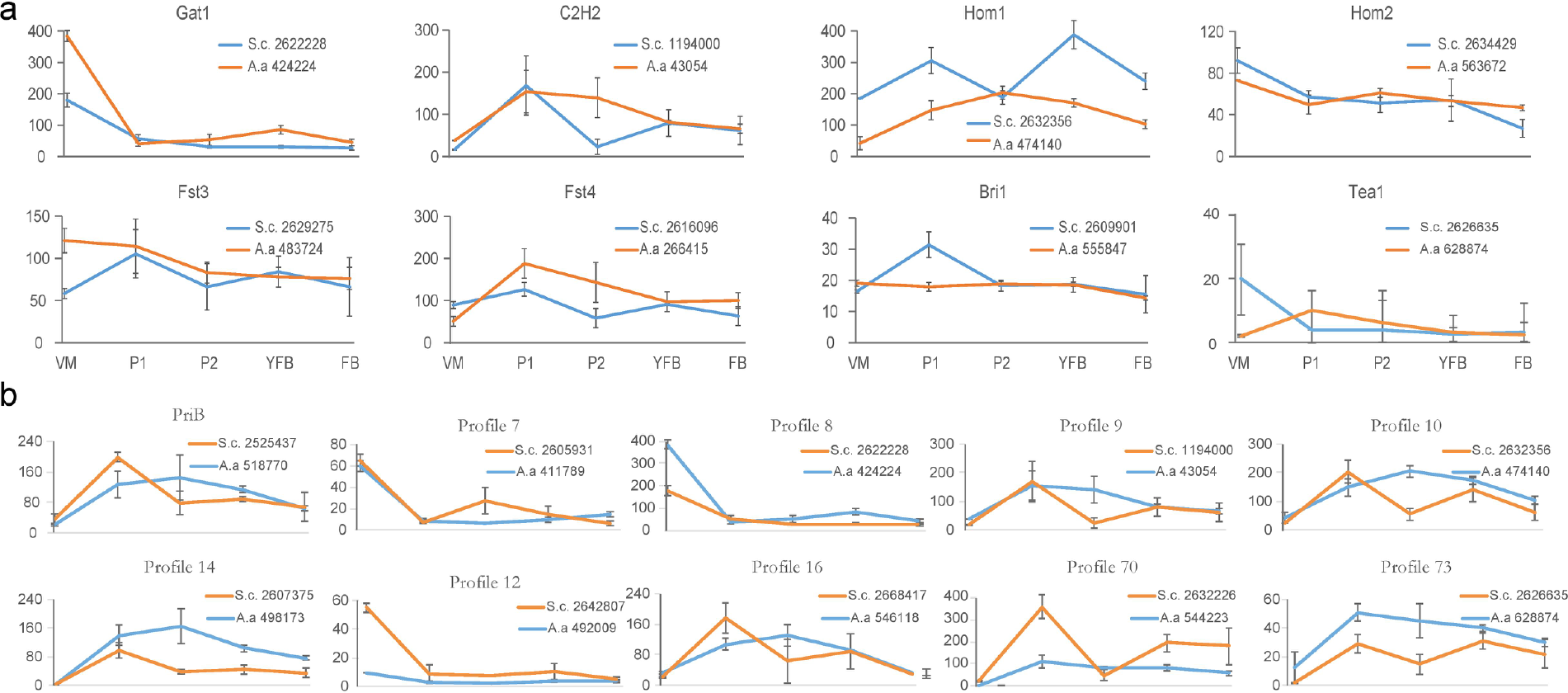
The expression patterns of developmentally regulated co-orthologous transcription factors and their similarity across the two species. A, Expression patterns for 8 previously characterized TFs in *S. commune* and *A. ampla*. B, developmentally regulated co-orthologous TFs in the two species with high expression dynamics during fruiting body development. *S. commune* and *A. ampla* genes are shown by blue and orange lines respectively.

Of the 252 co-orthologous TFs, 42 were developmentally regulated in both species, 27 of which had similar expression profiles between *A. ampla* and *S. commune*. Nine of the most interesting of these TFs are shown on Figure 5. Three of these genes showed highest expression in vegetative mycelia and are probably not relevant to fruiting body development. For the other six genes an upregulation was observed at the transition from vegetative mycelia to stage 1 primordia, which is compatible with potential roles in the initiation of fruiting body development or accompanying morphogenetic changes. Such TFs, with conserved, developmentally dynamic expression might be related to sculpting the specialized, cyphelloids fruiting body morphologies of *Auriculariopsis* and *Schizophyllum* or more widely conserved fruiting body functions. This also shows the power of comparative transcriptomics to identify genes with conserved expression patterns during fruiting body development ^83^ and to generate hypotheses that are testable by gene knockouts or functional assays.

### Small secreted proteins show dynamic expression in fruiting bodies

We detected several genes encoding short proteins with extracellular secretion signals (SSPs) in the fruiting body transcriptomes. In *A. ampla* and *S. commune* 316 and 354 SSPs were detected, respectively, half of which contained no known InterPro domains (Figure 6a, Supplementary Table 6). The SSPs in the two species belonged to 283 orthologs in *A. ampla*, and 315 in *S. commune*. Out of these, 133 orthologs were shared by the two species, whereas 150 and 182 were specific to *A. ampla* and *S. commune*, respectively (Figure 6b). The 133 shared orthologs contained 162 proteins, of which 39 were developmentally regulated in *A. ampla* and 158, with 54 developmentally regulated in *S. commune*. From these, 20 orthologs were developmentally regulated in both species (Figure 6d) and had a similar expression profile. Annotated SSPs in the two species (Figure 6c) had similar expression dynamics and mainly comprised hydrophobins, ceratoplatanins, CFEM domain containing proteins, concanavalin type lectins, and glycosyl hydrolases ^3^. The 150 species specific orthogroups in *A. ampla* contained 154 proteins, of which 32 are developmentally regulated. The 182 orthogroups in *S. commune* contained 196 proteins, of which 58 are developmentally regulated (Supplementary Figure 4).

**Figure 6.**
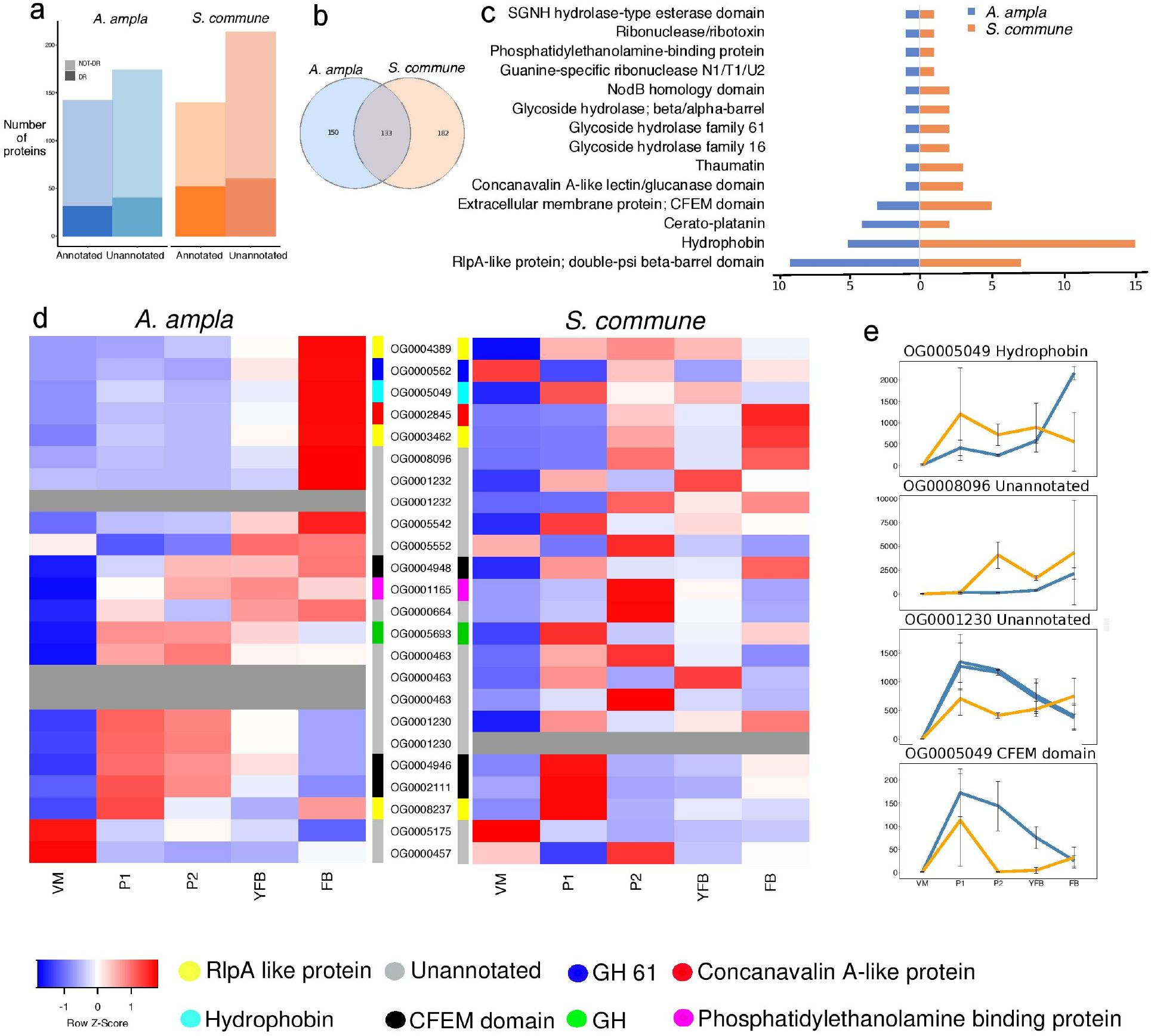
Small secreted proteins in fruiting body transcriptomes. A, The repertoires of annotated vs unannotated and developmentally regulated (DR) vs. non-developmentally regulated (NOT-DR) SSPs in fruiting body transcriptomes of *A. ampla* and *S. commune*. B, Venn-diagram depicting orthology relationships among SSPs of the two species. Number in each cell represent the number of shared or species-specific orthogroups. C, functional annotation terms (InterPro domains) present in SSPs of both *A. ampla* and *S. commune*. Terms specific to either species are not shown. D, expression heatmaps of co-orthologous SSPs in the two species. Orthogroup IDs are shown next to rows. Blue and red correspond to low and high expression, respectively. Greyed-out rows denote missing genes in orthogroup in which the 2 species did not have the same number of genes. Color coded bar next to heatmap shows functional annotations of the orthogroups. See Supplementary Figure 4 for heatmaps of species-specific genes. E, expression profiles of genes in four of the orthogroups through development, including two orthogroups of unannotated genes. Blue and orange denote *Auriculariopsis* and *Schizophyllum* genes, respectively. Variances across the three biological replicates are shown at corresponding developmental stages. See Supplementary Table 6 for protein IDs.

We detected several developmentally regulated SSP-s with no annotations, some of which showed high expression dynamics (FC>50, Figure 6e). We found 8, 15 and 2, *Auriculariopsis*-specific, *Schizophyllum*-specific and shared SSPs, respectively, with no known domains but a high expression dynamics (Supplementary Figure 4, Supplementary Table 6). For example, one of the orthogroups (Auramp1_494084, Auramp1_549528, Schco3_2664662) showed a considerable upregulation in stage 1 primordia in both species (FC=1166 - 1870), suggesting a role in the transition from vegetative mycelium to fruiting body initials. Such SSPs resemble effector proteins involved in cell-to-cell communication in ectomycorrhizal and pathogenic interactions between fungi and plants ^84,85^. Their expression in fruiting bodies raises the possibility that they play signaling roles and may be responsible for sculpting the fruiting bodies of these fungi. SSPs with an upregulation in morphogenetic processes (ECM root tips and/or fruiting bodies) have been reported in *Laccaria* ^21,84,86^ and *Pleurotus* ^87^ suggesting a role in tissue differentiation and that some of the SSPs initially found in ECM root tips may actually be morphogenetic in nature. Whether morphogenesis-related SSPs occur ubiquitously among mushroom-forming fungi and what is the mechanistic basis of their role, needs further research.

## Conclusions

In this study we presented the genome sequence of *Auriculariopsis ampla* and performed comparative genomic and transcriptomic analyses with *Schizophyllum commune* and other Agaricomycetes, to identify conserved genes related to fruiting body development and wood-decay. Our results showed that the two analyzed members of the Schizophyllaceae have a potential to shape our understanding of Agaricomycete biology. The CAZyme composition of *A. ampla* and *S. commune* compared to other 29 species suggests that the wood degrading strategies of the two species show similarity to WRs when concerned with cellulases, hemicellulases, and pectinolytic gene families. What sets them apart from WRs is the absence of class II PODs, which is also the case for BRs and ectomycorrhizal fungi. However, the reduction in ligninolytic genes is compensated by the presence of suberinases and tannases required to depolymerize important components bark, to which these species might be adapted.

Our analyses revealed a large number of genes with developmentally dynamic expression in fruiting bodies of both *A. ampla* and *S. commune*, including transcription factors (including 9 new conserved TFs), carbohydrate-active enzymes, heat shock proteins, aspartic proteases, as well as small secreted proteins. Particularly interesting are SSP-s with a highly dynamic expression through development, due to the role of SSPs in intracellular communication in pathogenic and ectomycorrhizal associations ^84,85^. Although mechanistic evidence is still lacking, it is conceivable that SSP-s with fruiting body specific expression might be involved in intercellular communication in fruiting bodies, similarly to their mycorrhiza and pathogenicity-related counterparts. This hypothesis would provide an explanation for the rich SSPs content of fruiting-body forming Agaricomycetes that are neither ectomycorrhizal or pathogenic ^3,84^.

Our data also suggest that despite the close phylogenetic relatedness of *Auriculariopsis* and *Schizophyllum*, their developmental transcriptomes have diverged significantly since their common ancestors, indicating a high rate of developmental gene expression in these taxa. Such divergence might be related to morphogenetic differences between the two species: while *A. ampla* produces simple cyphelloid (cup-shaped) fruiting bodies, those of *S. commune* consist of several congregated cyphelloid modules. Despite this divergence, several genes with a matching expression profile could be identified, highlighting conserved roles that await further experimentation. These data have the potential to highlight not only the genes involved in the development of cyphelloid fruiting bodies, but also that of other agaricomycete fruiting body types and as such, should be immensely useful to understanding the general principles and shared properties of fruiting body development in mushroom-forming fungi.

## Acknowledgements

This work was supported by the ‘Momentum’ program of the Hungarian Academy of Sciences (Contract No. LP2014/12 to L.G.N.) and the European Research Council (Grant No. 758161 to L.G.N.). The work conducted by the U.S. Department of Energy Joint Genome Institute (JGI), a DOE Office of Science User Facility, was supported by the Office of Science of the U.S. Department of Energy under Contract No. DE-AC02-05CH11231. This work was supported by the National Research, Development and Innovation office (Contract No. Ginop-2.3.2-15-00002, to L.G.N.).

## Methods

### Genome sequencing

The sequenced strain of *Auriculariopsis ampla* was collected as fruiting bodies on the bark of in Szeged, Hungary and cultured in liquid malt-extract medium (deposited in Szeged Microbiological Collections, under NL-1724). DNA was extracted using the DNeasy Blood & Tissue Culture kit (Qiagen), following the manufacturer’s protocol. The genome was sequenced using Pacific Biosciences RS II platform. Unamplified libraries were generated using Pacific Biosciences standard template preparation protocol for creating >10kb libraries. 5 ug of gDNA was used to generate each library and the DNA was sheared using Covaris g-Tubes (TM) to generate sheared fragments of >10kb in length. The sheared DNA fragments were then prepared using Pacific Biosciences SMRTbell template preparation kit, where the fragments were treated with DNA damage repair, had their ends repaired so that they were blunt-ended, and 5’ phosphorylated. Pacific Biosciences hairpin adapters were then ligated to the fragments to create the SMRTbell template for sequencing. The SMRTbell templates were then purified using exonuclease treatments and size-selected using AMPure PB beads. Sequencing primer was then annealed to the SMRTbell templates and Version P6 sequencing polymerase was bound to them. The prepared SMRTbell template libraries were then sequenced on a Pacific Biosciences RSII sequencer using Version C4 chemistry and 4-hour sequencing movie run times. Filtered subread data was assembled together with Falcon version 0.4.2 (https://github.com/PacificBiosciences/FALCON) and subsequently polished with Quiver version smrtanalysis_2.3.0. 140936.p5 (https://github.com/PacificBiosciences/GenomicConsensus).

For transcriptome, Stranded cDNA libraries were generated using the Illumina Truseq Stranded RNA LT kit. mRNA was purified from 1 ug of total RNA using magnetic beads containing poly-T oligos. mRNA was fragmented and reversed transcribed using random hexamers and SSII (Invitrogen) followed by second strand synthesis. The fragmented cDNA was treated with end-pair, A-tailing, adapter ligation, and 8 cycles of PCR. The prepared library was quantified using KAPA Biosystem’s next-generation sequencing library qPCR kit and run on a Roche LightCycler 480 real-time PCR instrument. The quantified library was then multiplexed with other libraries, and the pool of libraries prepared for sequencing on the Illumina HiSeq sequencing platform utilizing a TruSeq paired-end cluster kit, v4, and Illumina’s cBot instrument to generate a clustered flow cell for sequencing. Sequencing of the flow cell was performed on the Illumina HiSeq2500 sequencer using HiSeq TruSeq SBS sequencing kits, v4, following a 2×150 indexed run recipe. Illumina fastq files were QC filtered for artifact/process contamination and de novo assembled with Trinity v2.1.1 ^88^ and used for genome annotation

The genome was annotated using the JGI Annotation pipeline ^89^ and made available via JGI fungal genome portal MycoCosm (jgi.doe.gov/fungi;^89^). The data also deposited at DDBJ/EMBL/GenBank under the accession (*TO BE PROVIDED UPON PUBLICATION*).

### Fruiting protocol, RNA extraction and transcriptome sequencing Fruiting and RNA extraction

*Auriculariopsis ampla* was grown on sterilized poplar (*Populus alba*) bark and wood plugged into malt-extract medium in 250 ml glass beakers. Cultures were pre-incubated for 14 days in the dark at 30°C, then transferred to room temperature 60 cm under a light panel of 6 Sylvania Activa 172 Daylight tubes, with a 12 hr light/dark cycle and >90% relative humidity. Primordia started to develop 7 days after the transfer to light.

Vegetative mycelium, Stage 1 and 2 primordia, young and mature fruiting bodies were collected with sterilized forceps, flash-frozen in liquid nitrogen and stored at −80°C. Stage 1 and 2 primordia were defined as 0.1-1 mm closed, globular structures and 1-2 mm long initials with a central externally visible pit, respectively. We considered already opened fruiting bodies with a cup shape, 3-6 mm wide as young fruiting bodies and fully open ones as mature fruiting bodies. Total RNA was extracted from tissues ground to a fine powder in a mortar, using the Quick-RNA Miniprep kit (Zymo Research), following the manufacturer’s protocol. RNA samples with RIN>8 was saved for RNA-Sequencing. For each sample type three biological replicates were processed.

### RNA-Seq

Whole transcriptome sequencing was performed using the TrueSeq RNA Library Preparation Kit v2 (Illumina) according to the manufacturer’s instructions. RNA quality and quantity were assessed using RNA ScreenTape and Reagents on TapeStation (all from Agilent) and Qubit (ThermoFisher); only high quality (RIN >8.0) total RNA samples were processed. Next, RNA was DNaseI (ThermoFisher) treated and the mRNA was purified based on PolyA selection and fragmented. First strand cDNA synthesis was performed using SuperScript II (ThermoFisher) followed by second strand cDNA synthesis, end repair, 3’-end adenylation, adapter ligation and PCR amplification. Purification steps were performed using AmPureXP Beads (Backman Coulter). Final libraries were quality checked using TapeStation. Concentration of each library was determined using either the qPCR Quantification Kit for Illumina (Agilent) or the KAPA Library Quantification Kit for Illumina (KAPA Biosystems). Sequencing was performed on Illumina instruments using the HiSeq SBS Kit v4 250 cycles kit (Illumina) generating >20 million clusters for each sample.

### Bioinformatic analyses of RNA-Seq data

RNA-Seq analyses were carried out as reported earlier ^3,4^. Paired-end Illumina (HiSeq) reads were quality trimmed using CLC Genomics Workbench 9.5.2 (CLC Bio/Qiagen), removing ambiguous nucleotides as well as any low quality read ends. Quality cutoff value (error probability) was set to 0.05, corresponding to a Phred score of 13. Trimmed reads containing at least 40 bases were mapped using the RNA-Seq Analysis 2.1 package in CLC requiring at least 80% sequence identity over at least 80% of the read lengths; strand specificity was omitted. Reads with less than 30 equally scoring mapping positions were mapped to all possible locations while reads with more than 30 potential mapping positions were considered as uninformative repeat reads and were removed from the analysis.

“Total gene read” RNA-Seq count data was imported from CLC into R 3.0.2. Genes were filtered based on their expression levels keeping only those features that were detected by at least five mapped reads in at least 25% of the samples included in the study. Subsequently, “calcNormFactors” from “edgeR” 3.4.2 ^90^ was used to perform data scaling based on the “trimmed mean of M-values” (TMM) method73. Log transformation was carried out by the “voom” function of the “limma” package 3.18.13 ^91^. Linear modeling, empirical Bayes moderation as well as the calculation of differentially expressed genes were carried out using “limma”. Genes showing at least four-fold gene expression change with an FDR value below 0.05 were considered as significantly differentially expressed. Multi-dimensional scaling (“plotMDS” function in edgeR) was applied to visually summarize gene expression profiles revealing similarities between samples.

Developmentally regulated genes were defined as genes showing an >4-fold change in expression through development. In comparisons of vegetative mycelia and stage 1 primordia, we only considered genes upregulated in primordia, to exclude genes that showed a highest expression in vegetative mycelium because those might be related to processes not relevant for fruiting body development (e.g. nutrient acquisition).

### Phylogenetic analysis

Single-copy orthogroups were identified in MCL clusters of the 31 Agaricomycetes and were aligned by the l-ins-i algorithm of MAFFT ^92^. Ambiguously aligned regions were removed using the ‘strict’ settings of Trim-Al. Trimmed alignments longer than 100 amino acids were concatenated into a supermatrix. A maximum likelihood phylogenetic analysis was performed in RAxML 8.2.11 under the PROTGAMMALG model, with a gamma-distributed rate heterogeneity and a partitioned model. A bootstrap analysis in 100 replicates was performed.

### Identification of orthologous groups and their functional annotations

Groups of orthologous genes have been identified using OrthFinder v 1.1.8 ^93^, based on predicted protein sequences and the program's default parameters. Two analyses were performed, one to delimit orthogroups across 31 Agaricomycetes species and the second to find co-orthologs shared by *A. ampla* and *S. commune*, both using identical parameters. Functional annotation of proteins was carried out based on InterPro domains using InterProscan version 5.28-67.0 across the 31 fungal proteomes.

### Analyses of Carbohydrate Active Enzymes (CAZymes)

The CAZymes in the 31 species used in this study were annotated using the CAZy annotation pipeline ^94^. Copy numbers for most species were extracted from JGI Mycocosm, whereas those of *Flammulina velutipes, Coprinopsis marcescibilis*, and *Galerina marginata* were annotated for this study. Of all the families found in the dataset, we took into account the CAZyme families reported having a putative role in plant cell wall degradation ^6,8,9,22^(Supplementary Table 5) and analyzed their copy numbers across the 31 species. We used 45 CAZy families in our dataset (Supplementary Table 5) and divided them based on the degradation of celluloses, hemicelluloses, lignin, pectin.

In addition to these substrates, we also assessed genes encoding proteins with putative roles in suberin and tannin degradation. We extracted the best BLAST hits (BLAST 2.7.1+, e-value 0.001) from the 31 species for the homologs of proteins suggested to be related to suberin ^38,39^ and tannin degradation ^40^. We then identified the orthoMCL clusters of the 31 species containing the best hits. The proteins belonging to these clusters were used for further analysis as putative suberinases or tannases (Supplementary Table 5).

In order to get insights into the nutritional strategies employed by *A. ampla* and *S. commune*, we compared copy numbers of these species to that of 29 Agaricomycetes species with one of five known nutritional modes: brown rotters (BR), ectomycorrhizal (ECM), saprotrophs/litter decomposers/organic matter degraders (S/L/O), white rotters (WR) and uncertain. To analyze the grouping of the species according to their substrate degradation capabilities, phylogenetic PCA was performed using the phyl.pca ^95^ function from the R package phytools ^96^. A matrix of gene number normalized by proteome size in each organism (Supplementary Table 5), and the ML species tree, were used as input. Independent contrasts were calculated under the Brownian motion model and the parameter mode=“cov”.

### Analyses of transcriptome similarity

Pairwise comparisons of *A. ampla* and *S. commune* transcripts based on Pearson correlation coefficient among all replicates and developmental stages of 7369 OGs and among 1182 OGs containing at least one developmentally regulated gene were performed using custom Python script (pandas v 0.18.1 and Matplotlib v. 1.1.1 libraries). The same analysis has been performed for 252 single-copy ortholog transcription factors. The resulting matrix of Pearson correlation coefficients was plotted as a heatmap using the Matplotlib v. 1.1.1 pyplot framework. A scatterplot was constructed based on the log fold changes (FCs) of co-orthologs in *A. ampla* and *S. commune* using the ‘ggplot’ R package.

Heatmaps were created for developmentally regulated genes using the heatmap.2 function of R ‘gplot’ package. Hierarchical clustering with Euclidean distance calculation and averaged-linkage clustering was carried out on the FPKM values using ‘hclust’ function in R, and heatmaps was visualized using z-score normalization on the rows via the heatmap.2 function.

### Identification of transcription factors and other fruiting body genes

We identified transcription factor encoding genes in the proteomes of 31 Agaricomycetes species based on InterPro annotations. Only proteins containing domains with sequence-specific DNA binding ability were considered as transcription factors ^3^.

Carbohydrate active proteins were identified through the CAZy pipeline as described above. We extracted the kinases of the 2 species based on their InterPro domain composition and eliminated the ones involved in metabolism. Based on BLAST (*v*2.7.1+, e-value 0.001) against the classified kinome of *Coprinopsis cinerea* (Kinbase, Stajich et al., 2010), we assigned the best hits into the kinase categories of the query protein (^97,98^; www.kinase.com). Heatmaps were created using the Heatmap.2 function in R and are based on the FPKM values of developmentally regulated genes.

### Analyses of small secreted proteins

Small Secreted Proteins (SSPs) were identified for the two species to grasp species-specific and conserved SSPs in the two species. SSPs were defined as proteins shorter than 300 amino acids, having a signal peptide, an extracellular localization and no transmembrane domain. Proteins shorter than 300 amino acids were subjected to signal peptide prediction through SignalP 4.1^99^ with the option “eukaryotic”. The proteins having extracellular signal peptide were checked for their extracellular localization using WoLF PSORT 0.2 ^100^ with the option “fungi” and these were further checked for the absence of transmembrane helices, using TMHMM 2.0 ^101^.

---

**Supplementary Figure 1.**
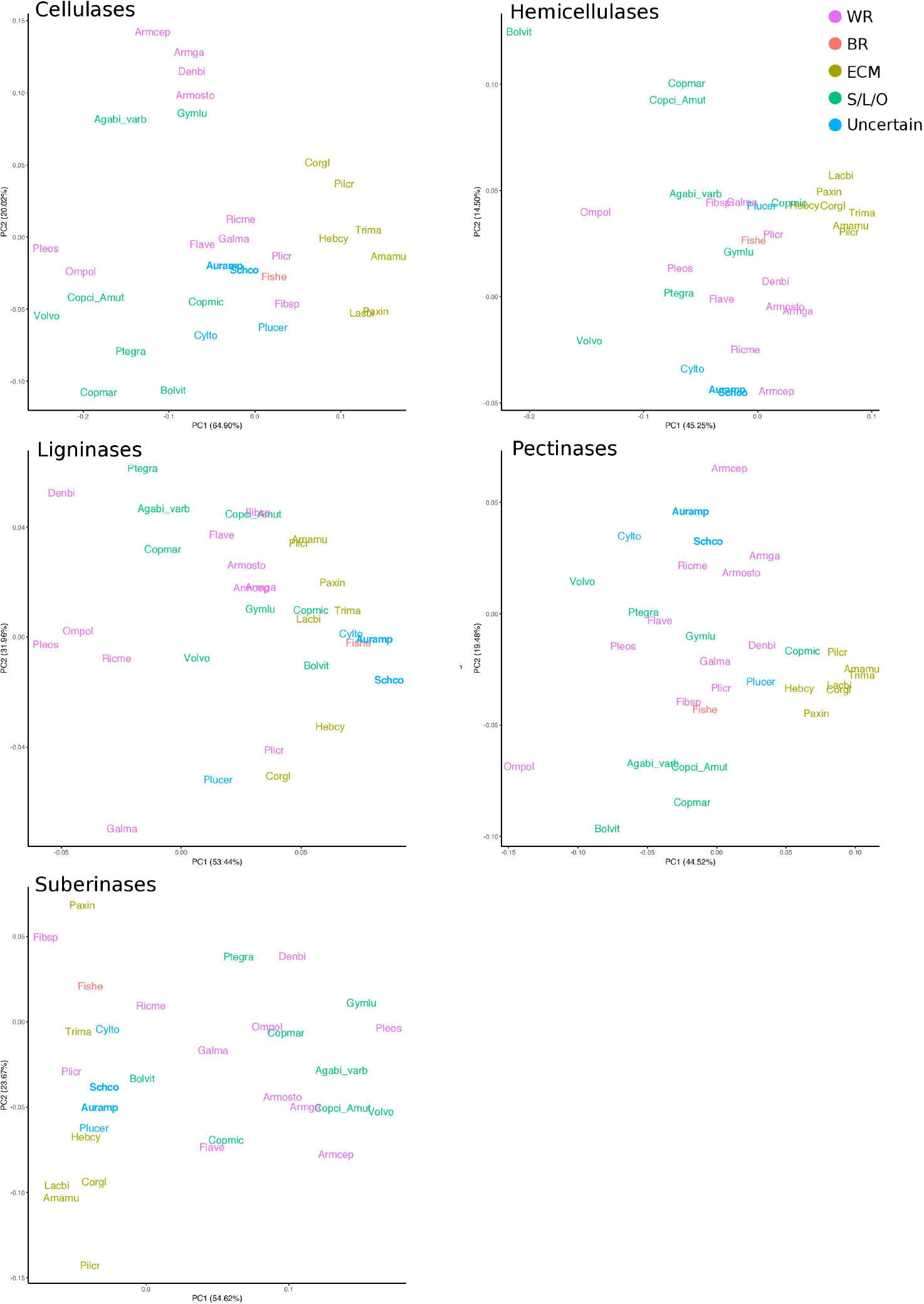
Phylogenetic Principal Component Analysis (PCA) based on CAZyme family copy numbers in 31 species. CAZymes categorized into 5 substrates - cellulases, hemicellulases, ligninases, pectinases and putative suberinases and the copy numbers normalized according to their proteome sizes along with ML species tree were used for the phylogenetic PCA. Species have been colored according to their nutritional modes.

**Supplementary Figure 2.**
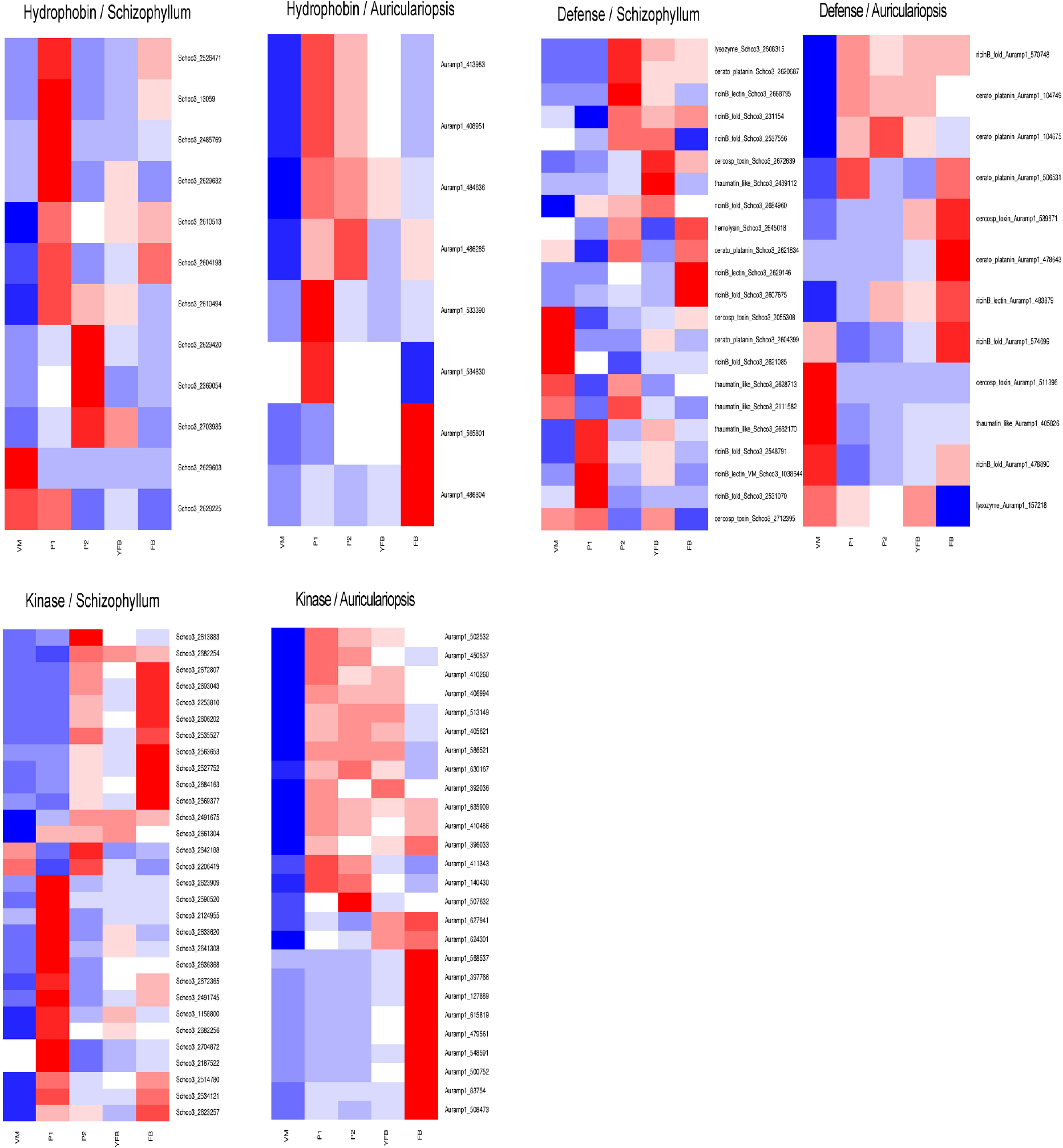
Heatmaps of developmentally regulated genes for families having a putative role in fruiting body formation. Developmental stages in both species are abbreviated as ‘VM’ vegetative mycelium; ‘P1’ stage 1 primordium; ‘P2’ stage 2 primordium; ‘YFB’ young fruiting body; and ‘FB’ fruiting body.

**Supplementary Figure 3.**
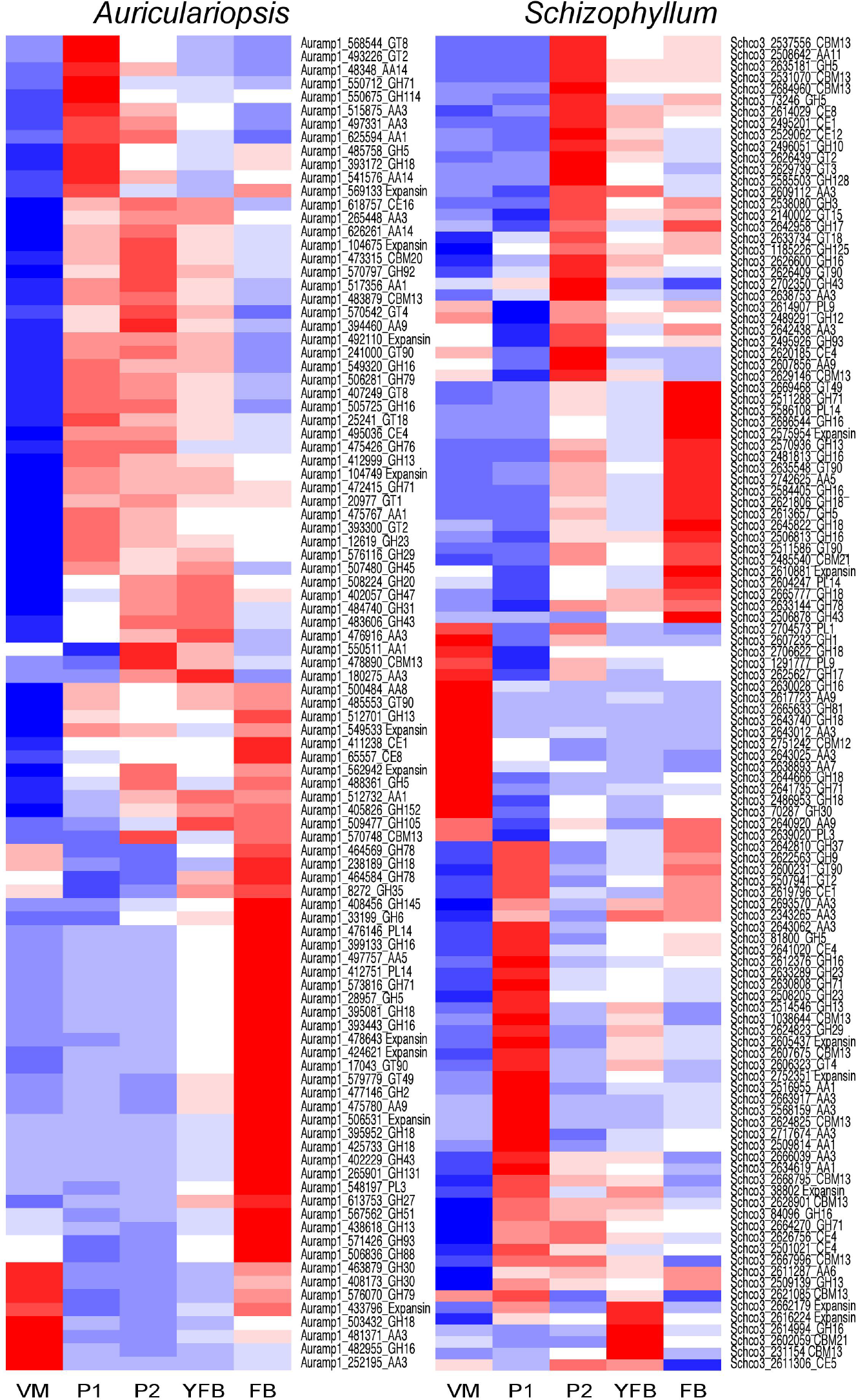
Heatmaps of developmentally regulated Carbohydrate Active Enzymes (CAZymes) in the two species. Abbreviations: ‘VM’ vegetative mycelium; ‘P1’ stage 1 primordium; ‘P2’ stage 2 primordium; ‘YFB’ young fruiting body; and ‘FB’ fruiting body.

**Supplementary Figure 4.**
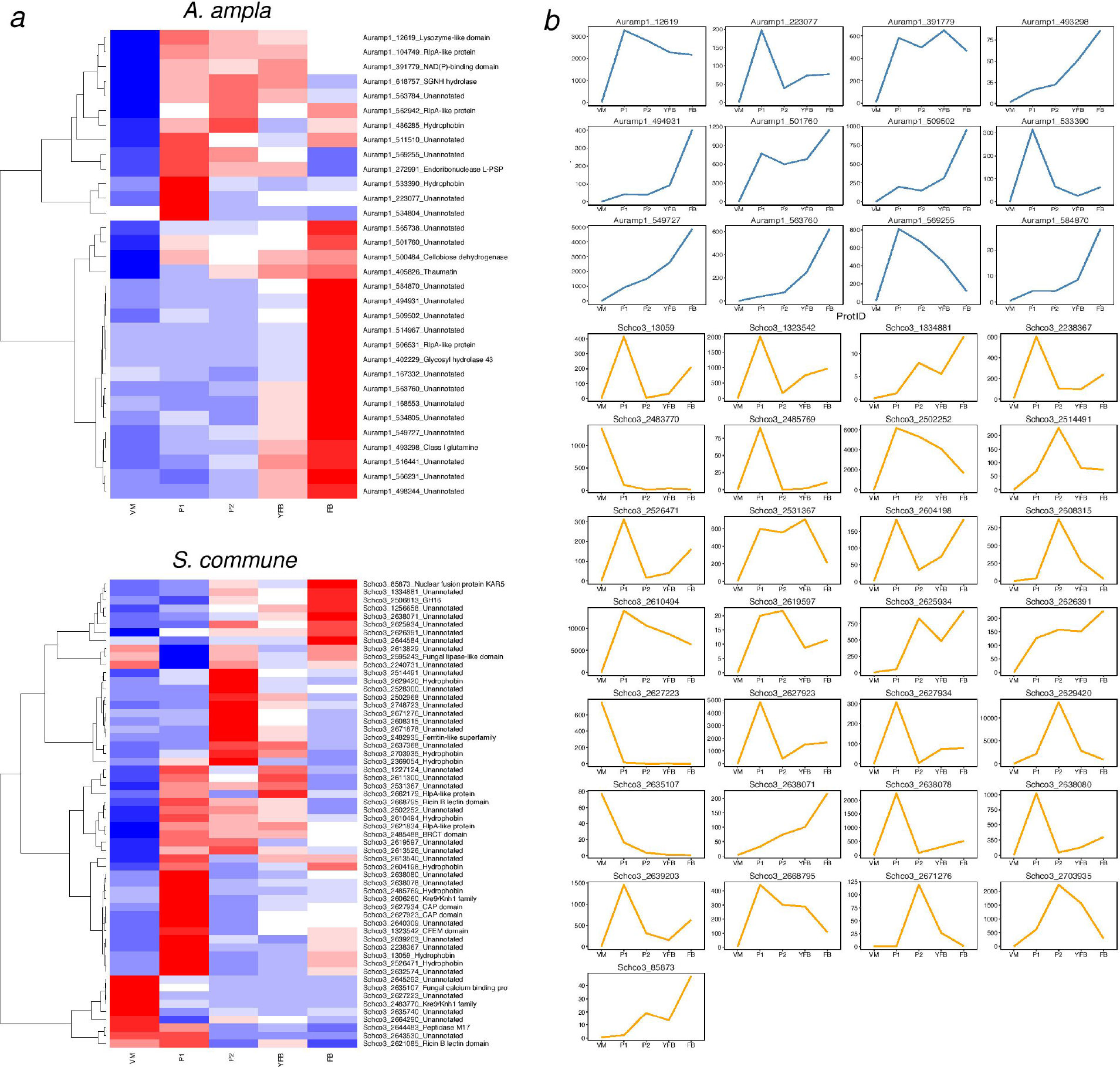
Developmentally regulated and species-specific small secreted proteins (SSPs) in the two species. A, Heatmaps of species specific SSPs in *A. ampla* (top) and *S. commune* (bottom); B, expression profiles of genes having high expression dynamics (FC>50). Genes in *A. ampla* and *S. commune* are shown in blue and orange respectively.

